# GtTR: Bayesian estimation of absolute tandem repeat copy number using sequence capture and high throughput sequencing

**DOI:** 10.1101/246108

**Authors:** Devika Ganesamoorthy, Minh Duc Cao, Tania Duarte, Wenhan Chen, Lachlan Coin

## Abstract

**Background:** Tandem repeats comprise significant proportion of the human genome including coding and regulatory regions. They are highly prone to repeat number variation and nucleotide mutation due to their repetitive and unstable nature, making them a major source of genomic variation between individuals. Despite recent advances in high throughput sequencing, analysis of tandem repeats in the context of complex diseases is still hindered by technical limitations.

**Methods:** We report a novel targeted sequencing approach, which allows simultaneous analysis of hundreds of repeats. We developed a Bayesian algorithm, namely – GtTR - which combines information from a reference long-read dataset with a short read counting approach to genotype tandem repeats at population scale. PCR sizing analysis was used for validation.

**Results:** We used a PacBio long-read sequenced sample to generate a reference tandem repeat genotype dataset with on average 13% absolute deviation from PCR sizing results. Using this reference dataset GtTR generated estimates of VNTR copy number with accuracy within 95% high posterior density (HPD) intervals of 68% and 83% for capture sequence data and 200X WGS data respectively, improving to 87% and 94% with use of a PCR reference. We show that the genotype resolution increases as a function of depth, such that the median 95% HPD interval lies within 25%, 14%, 12% and 8% of the its midpoint copy number value for 30X, 200X WGS, 395X and 800X capture sequence data respectively. We validated nine targets by PCR sizing analysis and genotype estimates from sequencing results correlated well with PCR results.

**Conclusions:** The novel genotyping approach described here presents a new cost-effective method to explore previously unrecognized class of repeat variation in GWAS studies of complex diseases at the population level. Further improvements in accuracy can be obtained by improving accuracy of the reference dataset.

## BACKGROUND

Repetitive DNA sequences make up almost half of the human genome [1]. A subset of these repeats are known as the tandem repeats (TRs) in which a stretch of DNA sequence (*i.e.* repeat unit) is located next to each other (*i.e.* in tandem). TRs with less than nine base pair repeat units are classified as microsatellites or short tandem repeats (STRs) and those with more than 10 base pair repeat units are known as minisatellites [2]. TRs, which have variable copy number in a population, are termed as variable number tandem repeats (VNTRs).

There are almost 1 million TRs in the human genome encompassing 4% of the entire genome [3], yet only few of these have been investigated in terms of disease association. Trinucleotide repeats in Fragile X Syndrome [4], Huntington’s disease [5], Spinobulbar muscular atrophy [6] and Spinocerebellar Ataxia [7] are few of the well-documented human diseases associated with TR variation. Notably these TRs are microsatellites; minisatelittes have not been well studied because of the limitations in analyzing longer length repeat units.

The contribution of genetic variation to complex There are almost 1 million TRs in plex disease susceptibility has been extensively studied in the recent years. Single nucleotide polymorphisms (SNPs) and copy number variations (CNVs) have been the major focus of large scale genome wide association studies (GWAS). However, VNTRs have been largely ignored in the context of complex diseases due to their sequence complexity. Historically, VNTRs have been considered as non-functional DNA, due to their repetitive and unstable nature. However, VNTRs are prone to high rates of copy number variation and mutation due to the repetitive unstable nature, which makes them a major source of genomic variation between individuals, which could potentially explain some of the phenotypic variation observed in complex diseases [8, 9].

Recent studies have shown that 10% to 20% of coding and regulatory regions contain VNTRs, suggesting that repeat variations could have phenotypic effects [10]. Association analysis have identified cis correlations of large tandem repeat variants with nearby gene expression and DNA methylation levels, indicating the functional effects of tandem repeat variations on nearby genomic sequences [11]. These findings show that TRs, which represent a highly variable fraction of the genome, can exert functionally significant effects. However, the possibility of exploring this collection of genetic variation is hindered by the difficulties in sequencing repetitive regions and the limitations of existing tools. As a result, the impact of tandem repeats on genomic variation between individuals as well as complex diseases remains largely unknown.

Traditionally, TR analysis has been carried out via restriction fragment length polymorphism (RFLP) analysis in which restriction enzymes are designed to fragment a target region, and genotyping was carried out by separation of fragments on a gel [12]. Recently, PCR amplification of the target loci, followed by capillary electrophoresis analysis was used to determine the fragment length of the alleles [13]. However, these techniques are only applicable to a specific target region and not scalable to high-throughput analysis. Hence, it limits the possibility of TR analysis in large-scale association studies.

Recent progress has been made in genotyping STRs using high-throughput short-read Illumina sequence data by use of local assembly techniques [14–16], which has led to insights into the role in variation in STR repeat length in controlling expression levels [17–19]. However, longer VNTRs remain intractable using these approaches with short to medium length reads. Sequencing reads which span the entire repeat regions could be informative to accurately genotype repeat copy number variation [20]. However this is not feasible for large scale analysis of longer TRs due to the high costs associated with long-read sequencing technologies.

We propose a novel genotyping approach with targeted capture sequencing, which can be used in combination with short read sequencing technologies to assess TR variation at a population scale. We first demonstrate that targeted sequence capture of repetitive TR regions are feasible. We describe a novel probabilistic algorithm (GtTR) for genotyping TRs from short read sequencing data (targeted capture sequencing or whole genome sequencing) by comparison of regional read-depth with a single long-read reference sample. Our analysis methodology requires the use of long read sequencing for only one sample to use as a reference, and can scale to population level with more economical short read sequencing technology. We demonstrate the accuracy of the estimates from GtTR by comparison with gold-standard PCR sizing analysis.

Our novel long read reference based genotyping approach of combining long read sequencing with targeted sequence capture using short read sequencing enables to genotype long TRs up to 5Kb in length and possibly longer with improved long read sequencing methods. It also provides a cost effective approach to genotype TRs for large scale analysis and has the potential to be applied in large scale genome wide association studies to uncover the genetic impact of long TRs on complex traits.

## METHODS

### Selection of Tandem Repeats for Analysis

This study was carried out as part of a larger study to investigate association between TRs and Obesity. The TRs targeted in this study were identified from SNP microarray intensity data collected on childhood obesity case control data (646 cases and 589 controls) and adult obesity case control data (709 cases and 197 controls) [21, 22]. Briefly, we selected microarray probes overlapping the VNTRs (as determined from the Tandem Repeats Database (TRDB) (4)) for association analysis. Principal component analysis (PCA) was used to identify association between obesity and the probe intensity measurements within the VNTR regions. A comparative analysis using Multiphen [23] was used to identify the top 50 VNTRs associated with obesity in each cohort (Child_Gender1, Child_Gender2 and Adult). We selected a combined total of 142 VNTRs for the targeted sequencing analysis. The selected TRs range from 112bp to 25236bp in length in the reference human genome and the number of repeat units range from 2 to 2300 repeats (Figure 1).

**Figure 1:**
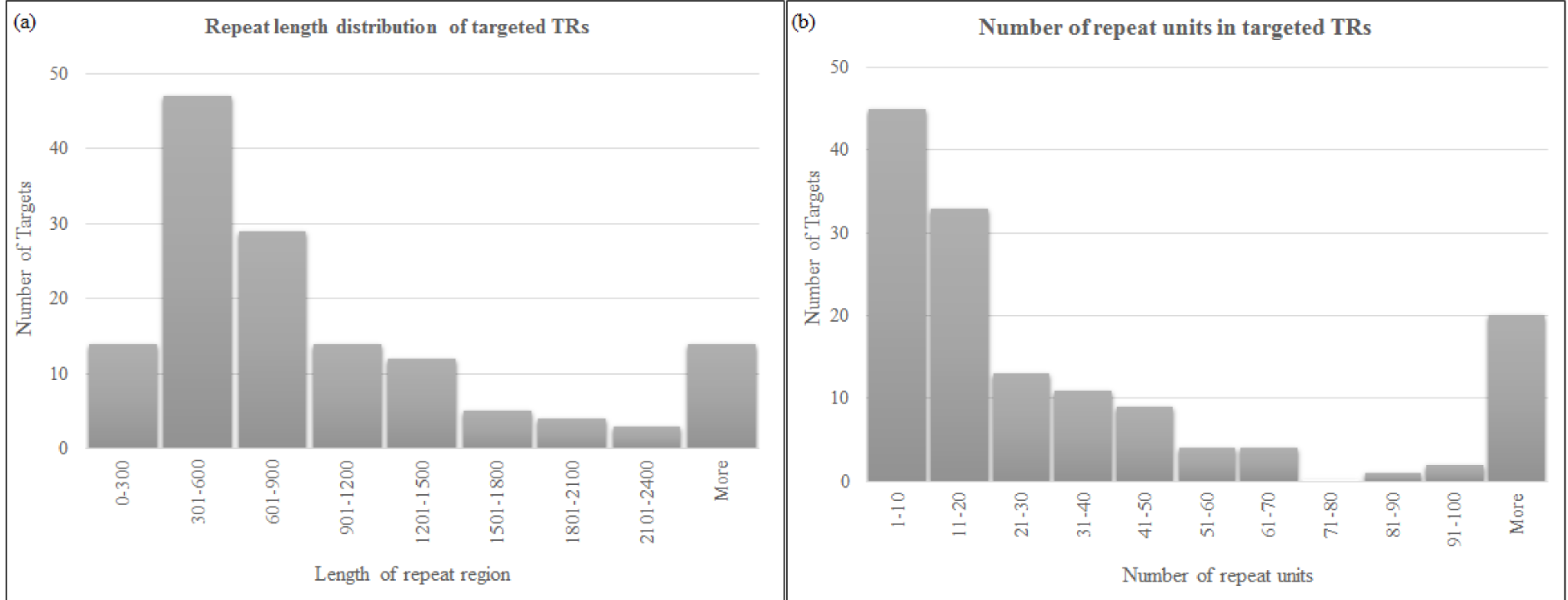
Distribution of (a) repeat length and (b) number of repeat units across the selected TRs.

### Probe design for selected TRs

Agilent SureSelect DNA design was used to design target probes to capture the targeted regions. The 142 selected VNTRs were used as targets for the design. 100bp flanking regions were also included as part of the target sequence. A high density tiling approach was used for probe design in target regions. Since the design is intended to target repeats, the repeat masker option was avoided to facilitate even probe coverage in the targeted region. Regions flanking the VNTRs were also included in the design. The size of the flanking region was determined by the size of the repeat region, at least 1000bp flanking sequence was included for each target. A high density tiling approach was used for probe design in flanking regions as well, however the repeat masker option was used to identify unique flanking sequences in the flanking region. Therefore the probe coverage in the flanking region is not evenly distributed.

### Samples for sequencing

DNA samples of CEPH/UTAH pedigree 1463 was purchased from Coriell Institute for Medical Research (USA). Seven family members from the pedigree was used for sequencing analysis and the samples encompass 3 trios (Trio 1 – NA12877, NA12878, NA12879; Trio 2 – NA12889, NA12890, NA12877; Trio 3 – NA12891, NA12892, NA12878).

### Capture and Illumina sequencing of targeted TRs

Seven samples were used for Illumina sequencing. Library preparation was performed using Agilent SureSelectXT Target Enrichment kit according to the manufacturer’s instructions. Briefly, DNA was fragmented to 600 to 800bp using micro-TUBE (Covaris). Fragments were end repaired, adapter ligated and amplified prior to target enrichment. Amplified fragments were hybridized to the designed capture probes for 24 hours. After hybridization, Streptavidin beads were used to capture the DNA fragments bound to the probes. Captured DNA was amplified using Illumina indexing adapters. Amplified libraries were sequenced on Illumina MiSeq with 300bp paired end sequencing. Samples NA12878 and NA12891 were sequenced as a pool of 4 samples in run 1 (other 2 samples in the pool are not included in the paper), whereas samples NA12877, NA12878, NA12879, NA12889, NA12890 and NA12892 were sequenced as a pool of 6 samples in run 2. Sample NA12878, which was used as reference sample was sequenced in both sequencing runs and data from sequencing run 1 was used as test sample and data from sequencing run 2 was used as reference sample.

### PCR analysis of VNTRs

PCR sizing analysis of VNTRs have inherent limitation due to repetitive sequences and size limitation of the PCR products for fragment analysis. Therefore only nine targeted VNTR regions which are less than 1Kb in repetitive sequence were validated by PCR sizing analysis in this study. Nevertheless, these nine regions include various repeat unit length and repeat sequence combinations to assess the accuracy of the genotypes determined from sequencing data. PCRs were performed using HotStar Taq DNA Polymerase (Qiagen) and PCR conditions were optimized for each PCR target. PCR products were purified and subjected to capillary electrophoresis on an ABI3500xL Genetic Analyzer (Applied BioSystems). Fragment sizes were analyzed using GeneMapper 4.0 (Applied BioSystems). Sanger sequencing was performed on PCR products to confirm the sequence of the repeat regions.

### Simulation of targeted sequencing data

We used simulated sequencing data to assess the accuracy of our genotyping algorithm. Generation of simulated data is described in Cao *et. al.* (2017) [24]. We first introduced SNPs and small indels to the reference human genome (hg19) to create 4 diploid genomes - Genome1, Genome2, Genome3 and Genome4. The rates for SNPs and indels were 2500 per MB and 280 per MB respectively, following the analysis of the 1000 genomes project [25]. We then introduced repeat variations into these genomes (Additional File 2 - Supplementary Table1). We simulated PacBio whole genome sequencing (WGS) data for Genome 2 and we simulated Illumina targeted sequencing data for these 142 loci for all 4 simulated genomes according to Cao et. al. (2017) [24]. For simulation of capture sequencing data, we sampled fragments from each genome according to the length distributions observed in real sequencing data (mean 800bp and standard deviation 100bp). We simulated approximately 1000X coverage for Illumina targeted sequencing data, however for downstream analysis we down-sampled to approximately 200X coverage to achieve comparable depth to the real targeted capture sequencing data.

**Table 1:** Genotype calls by GtTR using NA12878 Pacbio reference on targeted capture sequencing data compared with PCR sizing analysis.

| Sample | Method | Target |  |  |  |  |  |  |  |  | Correlation |
| --- | --- | --- | --- | --- | --- | --- | --- | --- | --- | --- | --- |
|  |  | VNTR_8 | VNTR_32 | VNTR_57 | VNTR_86 | VNTR_87 | VNTR_93 | VNTR_109 | VNTR_112 | VNTR_120 |  |
| NA12877 | PCR | 12.5 | 21.8 | 12.1 | 2.0 | 7.0 | 2.0 | 16.0 | 3.4 | 2.0 | 0.9762 |
|  | Capture_Seq | 11.5 | 28.4 | 13.5 | 2.7 | 6.1 | 2.1 | 16.2 | 3.7 | 3.4 |  |
|  | 95% HPD interval | (10.2-12.8) | (18.8-38) | (11.9-15.2) | (2.07-3.33) | (5.43-6.7) | (1.76-2.52) | (14.4-18) | (3-4.35) | (2.41-4.38) |  |
| NA12878<sup>#</sup> | PCR | 12.0 | 9.5 | 11.6 | 2.0 | 7.5 | 2.0 | 16.0 | 3.4 | 2.0 | 0.9930 |
|  | Capture_Seq | 12.1 | 9.9 | 13.5 | 2.6 | 7.8 | 2.3 | 17.6 | 3.8 | 3.5 |  |
|  | 95% HPD interval | (10.9-13.3) | (5.5-14.7) | (12.1-15) | (2.12-3.14) | (7.1-8.41) | (1.97-2.69) | (16-19.2) | (3.19-4.35) | (2.64-4.36) |  |
|  | *PacBio WGS\** | 12.0 | 11.9 | 11.6 | 2.5 | 7.5 | 2.0 | 16.3 | 3.5 | 3.2 |  |
| NA12879 | PCR | 12.5 | 9.8 | 10.5 | 2.0 | 8.5 | 2.0 | 16.0 | 2.9 | 2.0 | 0.9738 |
|  | Capture_Seq | 12.6 | 11.7 | 14.4 | 2.6 | 8.2 | 2.0 | 16.3 | 3.2 | 3.0 |  |
|  | 95% HPD interval | (11.3-14) | (6-17.4) | (12.8-16) | (2.09-3.18) | (7.44-8.92) | (1.65-2.32) | (14.6-17.9) | (2.68-3.8) | (2.12-3.81) |  |
| NA12889 | PCR | 12.0 | 22.3 | 15.6 | 2.0 | 8.0 | 2.0 | 16.9 | 3.9 | 2.0 | 0.9871 |
|  | Capture_Seq | 11.1 | 27.0 | 20.3 | 2.2 | 7.7 | 2.2 | 21.3 | 4.1 | 3.4 |  |
|  | 95% HPD interval | (9.97-12.3) | (18.5-35.5) | (18.3-22.4) | (1.73-2.63) | (7.01-8.36) | (1.8-2.5) | (19.4-23.2) | (3.45-4.68) | (2.55-4.33) |  |
| NA12890 | PCR | 12.5 | 9.7 | 10.5 | 2.0 | 7.5 | 2.0 | 15.1 | 2.9 | 2.0 | 0.9853 |
|  | Capture_Seq | 13.5 | 10.3 | 11.6 | 2.2 | 7.2 | 2.2 | 13.4 | 3.1 | 3.0 |  |
|  | 95% HPD interval | (12.1-14.9) | (5.01-15.6) | (10.2-13) | (1.79-2.72) | (6.52-7.87) | (1.85-2.58) | (12-14.8) | (2.52-3.62) | (2.13-3.85) |  |
| NA12891 | PCR | 12.5 | 9.3 | 10.6 | 2.0 | 7.0 | 2.0 | 15.1 | 2.9 | 2.0 | 0.9882 |
|  | Capture_Seq | 11.7 | 10.4 | 12.1 | 2.6 | 6.8 | 2.1 | 16.2 | 3.1 | 3.5 |  |
|  | 95% HPD interval | (10.6-12.8) | (5.87-14.9) | (10.8-13.3) | (2.14-3.08) | (6.23-7.36) | (1.79-2.42) | (14.8-17.6) | (2.61-3.52) | (2.65-4.25) |  |
| NA12892 | PCR | 12.5 | 9.8 | 12.6 | 2.0 | 9.0 | 2.0 | 16.9 | 3.9 | 2.0 | 0.9833 |
|  | Capture_Seq | 11.0 | 8.8 | 12.9 | 3.0 | 8.4 | 2.3 | 18.3 | 4.0 | 3.4 |  |
|  | 95% HPD interval | (9.85-12.2) | (3.88-13.8) | (11.4-14.3) | (2.36-3.58) | (7.63-9.1) | (1.93-2.7) | (16.5-20.1) | (3.33-4.6) | (2.43-4.27) |  |

### Public data used in the study

Illumina WGS data on CEPH Pedigree 1463 samples were downloaded from ENA with accession number PRJEB3381 and PRJEB3246 [26]. PacBio WGS data on NA12878 sample was downloaded from SRA with accession numbers SRX627421 and SRX638310 [27]

### Genotyping TRs from PacBio Sequencing data

PacBio WGS data on NA12878 sample was mapped to the whole genome hg19 reference using BLASR [28]. We used VNTRTyper (https://github.com/mdcao/japsa), an in-house tool to genotype TRs from long read PacBio sequencing data. Recently a similar tool - adVNTR was reported by Bakhtiari *et. al.* (2017) [29]. Briefly, VNTRTyper takes advantage of the long read sequencing to identify the number of repeat units in the TR regions. Firstly, the tool identifies reads that span the repeat region and applies Hidden Markov Models (HMM) to align the repetitive portion of each read to the repeat unit. Then it estimates the multiplicity of the repeat unit in a read using a profile HMM. A threshold of 2 supporting reads per genotype was used to estimate genotypes. Details of VNTRtyper analysis is provided in Additional file 1 – Supplementary information.

### Sequencing analysis of TRs from Illumina Sequencing data

Both Illumina targeted capture sequencing data and WGS data were mapped to the human genome hg19 reference using BWA-MEM [30]. We developed “GtTR”, which is a read-depth Bayesian model for estimating both the maximum a-posteriori repeat count genotype, as well as the standard error and 95% high posterior density (HPD) interval in this estimate. GtTR estimates a scaled relative repeat count of the test sample to a reference sample. Define R_rep_, R_flank_, S_rep_, S_flank_ as the read count for the repeat and flanking region reference (R) and the test sample (S) respectively.

We calculate a frequentist estimate of RCN as

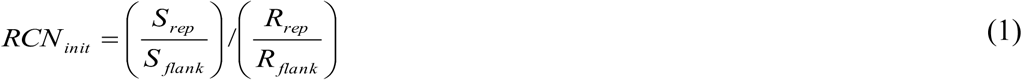

We calculate a posterior probability over a discretized set of possible RCN values as follows

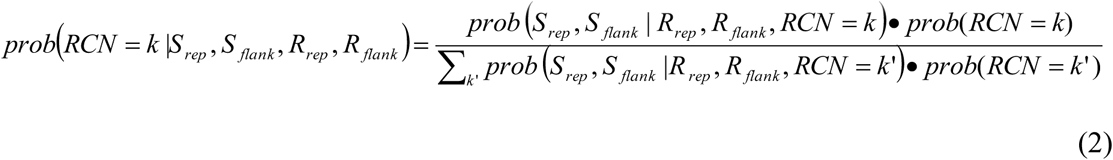

where

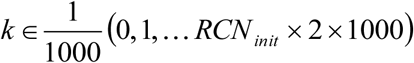

Assume that prob(RCN=k | R_rep_, R_flank_) = prob(RCN=k). We place a uniform prior on prob(RCN=k).

We model the expected number of reads in the repeat region (R_rep_), conditional on the total number of reads in the region (flanking plus repeats) using a beta-binomial distribution *prob S_rep_|S_flank_,R_flank_, RCN) ~ Binomial (S_rep_|number of trials S_flank_+S_rep_, p_succes_)*
with

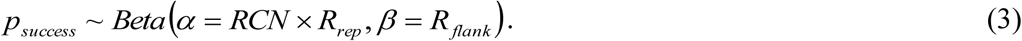

This is a model in which the proportion of reads expected to come from the repetitive region scales with RCN.

From equation 3 we calculate the maximum a-posteriori RCN (RCN_MAP_) as well as the smallest range which contains X% of the posterior probability mass (defined as the high posterior density interval), where the default for X is 95%. Finally, we rescale these values by multiplying by the reference genotype, as determined either by an estimate derived from PacBio data or PCR analysis. Details of GtTR (https://github.com/mdcao/japsa) analysis is provided in Additional File 1 – Supplementary information.

### Statistical Analysis

To calculate the accuracy as a function of HPD interval, we use GtTR to calculate the HPD interval for 10%, 20%, 30%, 40%, 50%, 60%, 70%, 80%, 90% and 95% of posterior probability mass respectively at all loci for which we have gold-standard PCR sizing results. We use the number of PCR sizing results which lie inside and outside these HPD intervals to estimate the accuracy, as well as 95% confidence intervals (CI) in this estimate (using the binom.confint function in the binom R package).

We also plot the distribution of half relative width (HRW) of the 95% HPD intervals. We calculate this value as 95% HRW-HPD = (HPD_upper – HPD_lower)/(HPD_upper + HPD_lower). For a non-skewed posterior distribution, we can interpret 95% HRW-HPD = x as the value x such that 95% of the posterior mass lies with HPD-midpoint +/- x * HPD-midpoint. We estimate this distribution over all captured TR regions for which we had sufficient long-read sequence coverage (122 regions). We also estimated this cumulative distribution after partitioning regions based on the average short read coverage depth in order to investigate the role of short-read sequence read depth on genotype resolution.

## RESULTS

We developed a novel approach to genotype tandem repeats from targeted capture sequencing which integrates a single whole-genome long-read reference sequence with short-read sequencing data. We evaluated this approach using a combination of simulated and real sequencing data.

### Using short read sequence to genotype TRs

The short-read length of Illumina sequencing reads (< 300bp) are not sufficient to span the entire repeat region and flanking region, which presents a hurdle for genotyping repeat regions. Here, we propose a novel algorithm ‘GtTR’ to utilize the cost-effective short-read sequencing method for genotyping repeat regions. We use a control sample with known genotype, which is determined from long-read sequencing as a reference to improve the accuracy of genotyping from short-read sequencing data (Figure 2). Due to the use of read-depth based approach genotypes determined from short-read sequencing data will be an average of the two alleles instead of the exact genotype of the two alleles.

**Figure 2:**
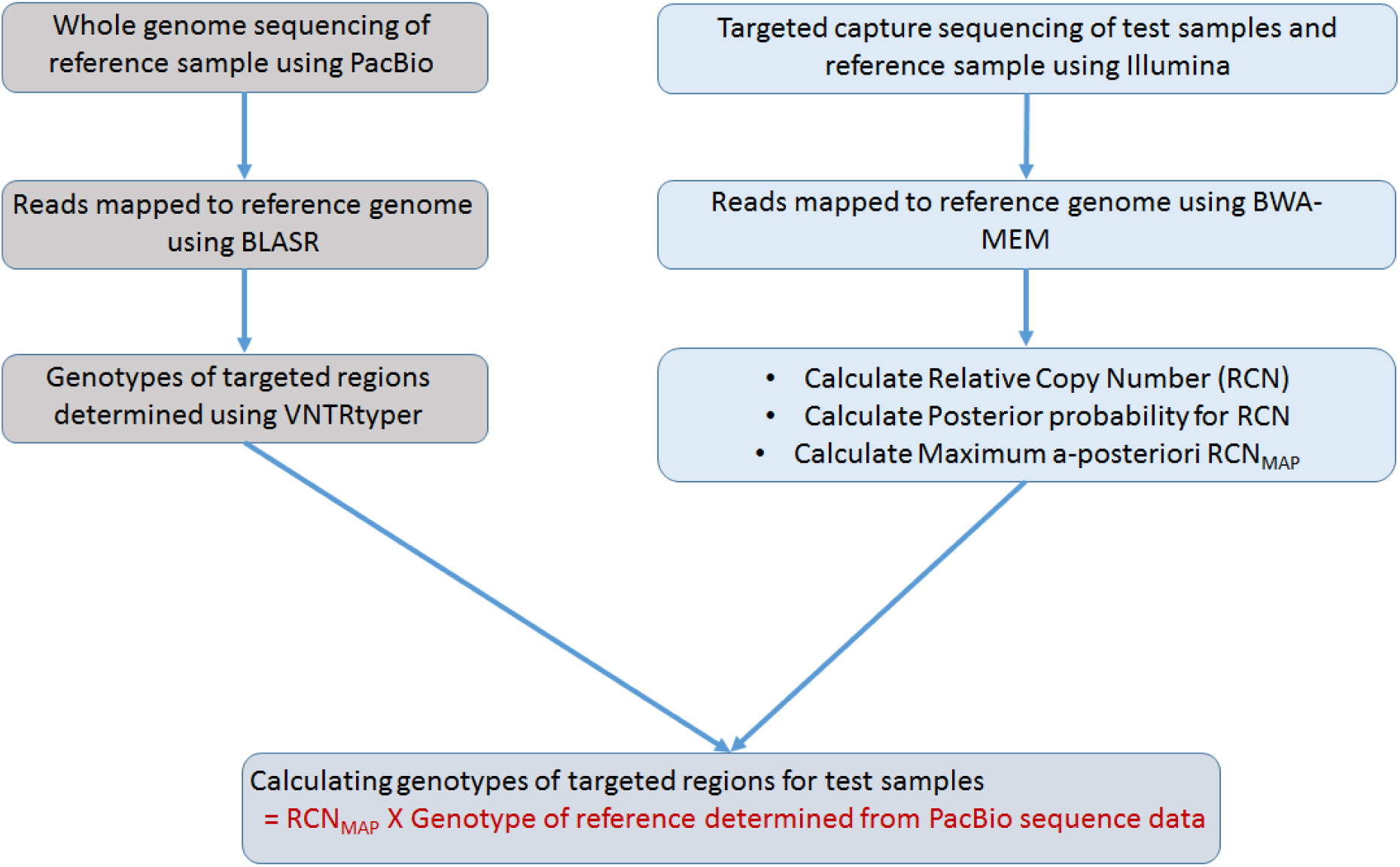
GtTR analysis pipeline for Illumina short-read sequencing data.

### Evaluating performance of GtTR using simulated data

We simulated Illumina targeted capture sequencing data from the targeted VNTR regions for Genome1, Genome2, Genome3 and Genome4 (see Methods). We also simulated PacBio WGS data from simulated Genome2, hence simulated Genome2 sample was used as the reference sample in the GtTR analysis pipeline. VNTRtyper was applied to simulated PacBio data on Genome2 to determine the genotypes of the targeted VNTR regions. PacBio WGS simulated data only had sufficient coverage for 119 targets to determine the genotype. VNTRtyper identified at least one allele correctly for 92 targets (Additional File 2 - Supplementary Table 2). The correlation values between simulated and observed genotypes were 0.9980, 0.9969 and 0.9971 for allele 1, allele 2 and both alleles, respectively, indicating that the VNTRtyper method produces an accurate estimation of the genotypes.

**Table 2:** Genotype estimates by GtTR on Illumina WGS data compared with PCR sizing analysis.

| Sample (Coverage) | Method | Target_ID |  |  |  |  |  |  |  |  | Correlation |
| --- | --- | --- | --- | --- | --- | --- | --- | --- | --- | --- | --- |
|  |  | VNTR_8 | VNTR_32 | VNTR_57 | VNTR_86 | VNTR_87 | VNTR_93 | VNTR_109 | VNTR_112 | VNTR_120 |  |
| NA12877 (30X) | PCR | **12.5** | 21.8 | 12.1 | 2.0 | 7.0 | 2.0 | 16.0 | 3.4 | 2.0 | 0.9827 |
|  | Illumina_WGS | 10.6 | 26.6 | 13.1 | 3.0 | 6.5 | 2.1 | 19.6 | 3.1 | 2.6 |  |
|  | 95% HPD interval | (7.94-13.2) | (16.5-36.8) | (9.99-16.2) | (2.12-3.95) | (4.88-8.03) | (1.47-2.8) | (15-24.2) | (1.89-4.29) | (0.784-4.46) |  |
| NA12877 (200X) | PCR | 12.5 | 21.8 | 12.1 | 2.0 | 7.0 | 2.0 | 16.0 | 3.4 | 2.0 | 0.9756 |
|  | Illumina_WGS | 12.4 | 29.2 | 13.2 | 2.5 | 7.0 | 2.3 | 15.5 | 3.2 | 3.0 |  |
|  | 95% HPD interval | (10.6-14.3) | (22.4-35.9) | (11.2-15.2) | (2-3.05) | (5.89-8.03) | (1.84-2.72) | (12.9-18.1) | (2.32-3.99) | (1.73-4.24) |  |
| NA12878 (30X) | PCR | 12.0 | 9.5 | 11.6 | 2.0 | 7.5 | 2.0 | 16.0 | 3.4 | 2.0 | 0.9251 |
|  | Illumina_WGS | 12.0 | 17.2 | 11.2 | 2.3 | 7.5 | 2.1 | 21.5 | 3.6 | 3.8 |  |
|  | 95% HPD interval | (9.24-14.8) | (8.72-25.6) | (8.24-14.1) | (1.51-3.06) | (5.74-9.25) | (1.42-2.76) | (16.7-26.4) | (2.18-5.11) | (1.51-6.16) |  |
|  | *PacBio* WGS* | 12.0 | 11.9 | 11.6 | 2.5 | 7.5 | 2.0 | 16.3 | 3.5 | 3.2 |  |
| NA12879 (30X) | PCR | 12.5 | 9.8 | 10.5 | 2.0 | 8.5 | 2.0 | 16.0 | 2.9 | 2.0 | 0.9659 |
|  | Illumina_WGS | 11.9 | 11.4 | 12.3 | 2.2 | 9.6 | 2.2 | 23.2 | 2.1 | 3.2 |  |
|  | 95% HPD interval | (9.06-14.7) | (4.59-18.1) | (9.26-15.4) | (1.48-2.96) | (7.53-11.7) | (1.51-2.79) | (18.2-28.2) | (1.08-3.04) | (1.08-5.31) |  |
| NA12889 (30X) | PCR | 12.0 | 22.3 | 15.6 | 2.0 | 8.0 | 2.0 | 16.9 | 3.9 | 2.0 | 0.9883 |
|  | Illumina_WGS | 11.1 | 25.3 | 18.1 | 2.6 | 7.7 | 2.2 | 21.4 | 3.8 | 3.3 |  |
|  | 95% HPD interval | (8.47-13.7) | (16-34.5) | (14.4-21.9) | (1.78-3.34) | (5.96-9.37) | (1.55-2.89) | (16.8-26) | (2.37-5.12) | (1.32-5.31) |  |
| NA12890 (30X) | PCR | 12.5 | 9.7 | 10.5 | 2.0 | 7.5 | 2.0 | 15.1 | 2.9 | 2.0 | 0.9640 |
|  | Illumina_WGS | 12.7 | 10.9 | 11.0 | 2.6 | 8.2 | 1.8 | 22.0 | 2.8 | 3.3 |  |
|  | 95% HPD interval | (9.66-15.8) | (3.96-17.9) | (7.92-14) | (1.72-3.51) | (6.34-10.1) | (1.15-2.47) | (16.9-27) | (1.56-4.02) | (1.05-5.48) |  |
| NA12891 (30X) | PCR | 12.5 | 9.3 | 10.6 | 2.0 | 7.0 | 2.0 | 15.1 | 2.9 | 2.0 | 0.9570 |
|  | Illumina_WGS | 12.6 | 9.8 | 12.6 | 2.5 | 7.4 | 2.1 | 23.0 | 2.2 | 3.5 |  |
|  | 95% HPD interval | (9.71-15.5) | (3.2-16.4) | (9.43-15.7) | (1.66-3.3) | (5.63-9.1) | (1.42-2.74) | (17.8-28.2) | (1.17-3.27) | (1.29-5.72) |  |
| NA12892 (30X) | PCR | 12.5 | 9.8 | 12.6 | 2.0 | 9.0 | 2.0 | 16.9 | 3.9 | 2.0 | 0.9704 |
|  | Illumina_WGS | 12.4 | 11.4 | 15.6 | 2.6 | 9.0 | 2.1 | 24.6 | 3.1 | 2.6 |  |
|  | 95% HPD interval | (9.64-15.2) | (4.83-18) | (12.2-18.9) | (1.78-3.44) | (7.12-11) | (1.39-2.8) | (19.5-29.8) | (1.88-4.29) | (0.691-4.45) |  |

\*Genotype estimates from PacBio WGS on NA12878 sample included for comparison.

The GtTR analysis pipeline was applied to all 4 simulated Illumina targeted sequencing data set to determine the repeat count genotypes, as well as the relative standard error in the estimate of the genotypes at the targeted VNTR regions. Genotype estimates from GtTR were compared with the simulated genotypes for all 4 simulated data sets (Figure 3, Additional File 2 - Supplementary Table 3).

**Figure 3:**
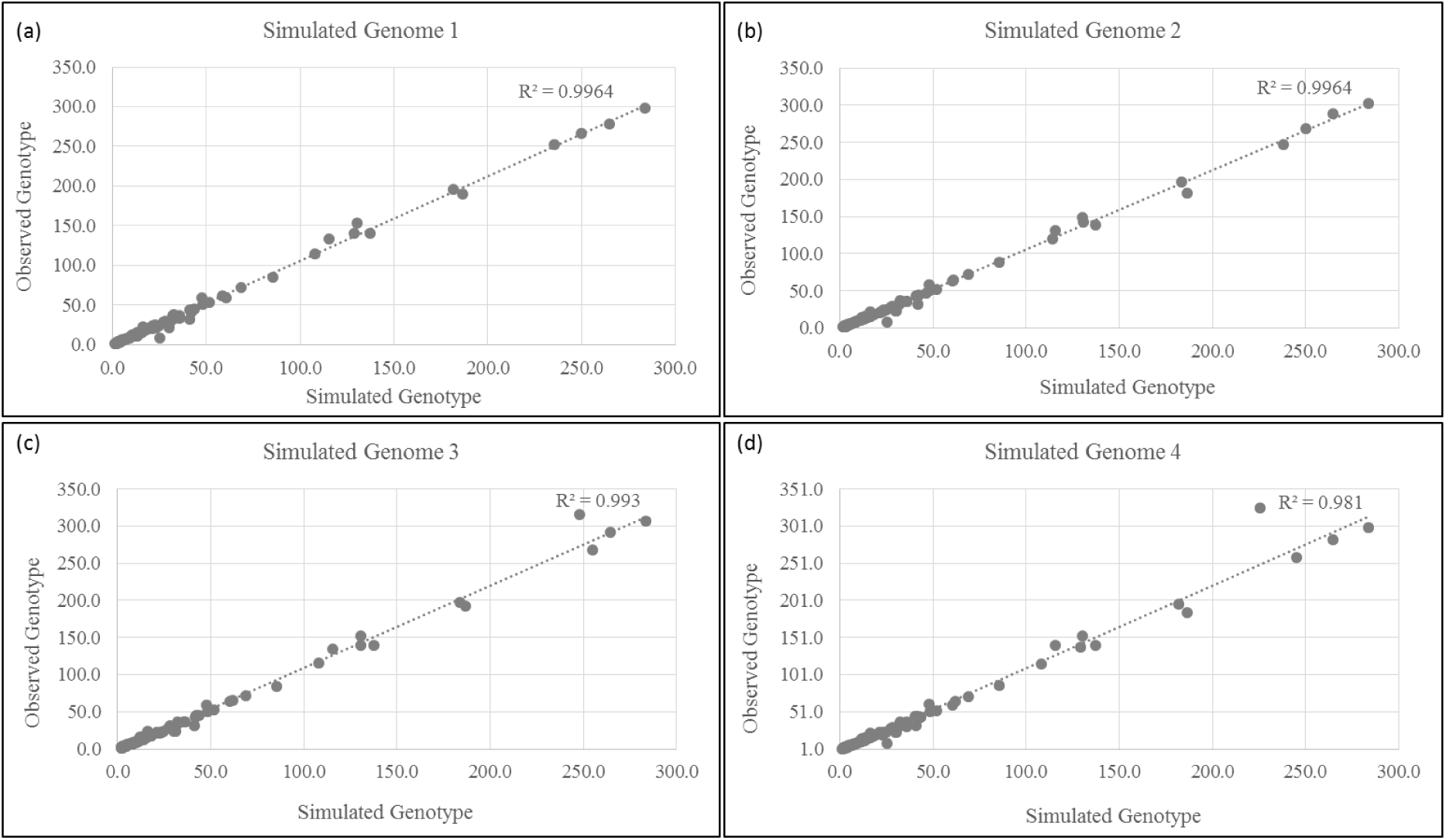
Correlation between GtTR genotype estimates and simulated genotypes on 4 simulated Illumina targeted sequencing datasets.

GtTR estimated genotypes were 96.6% accurate (CI: 94.6% - 98.0%) within 95% HPD intervals (Figure 4a). The median half relative width of 95% HPD interval was 11.2% (based on median depth of coverage across 4 samples and 119 targets of 578X coverage, Figure 4b)).

Half relative width of 95% HPD interval decreased to 8.8% amongst loci with depth between 800X and 1000X (Figure 4c).

**Figure 4:**
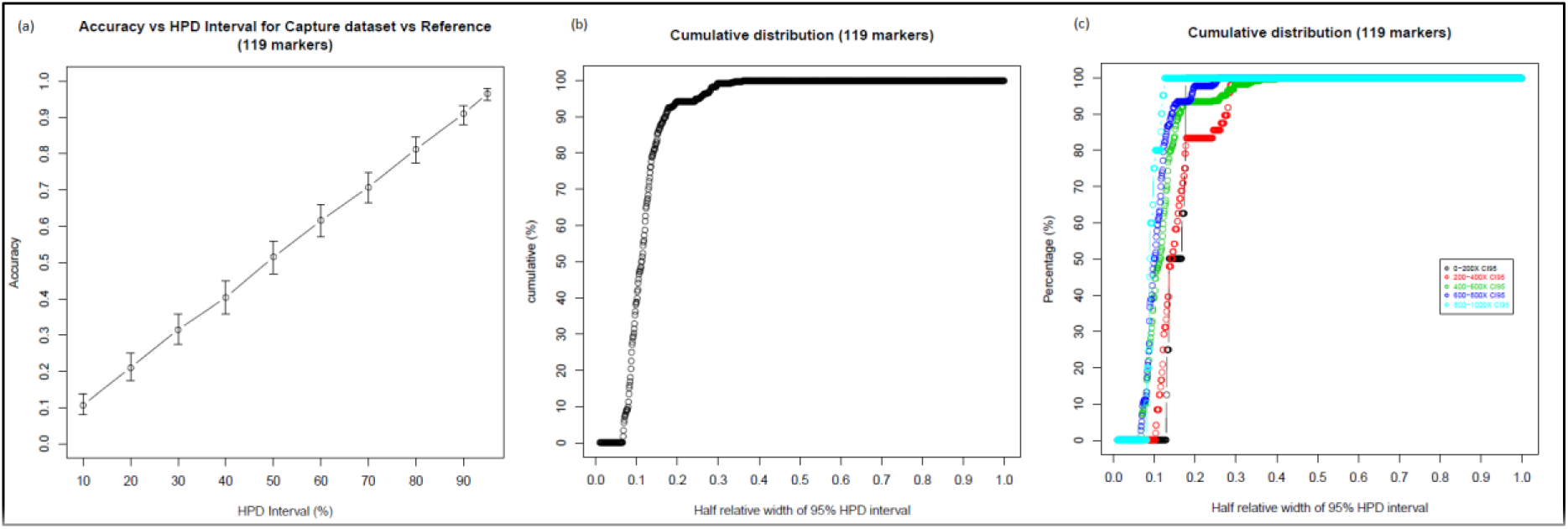
(a) Accuracy of the genotype estimates by GtTR at varying HPD intervals for simulated capture data using genotypes from long read sequence data as a reference genotype. Error bars represent 95% binomial confidence intervals. (b) Overall cumulative distribution of half relative width of 95% HPD interval in genotype estimates (c) Cumulative distribution of half relative width of 95% HPD interval in genotype estimates stratified by sequence coverage.

### Developing a global reference sample for GtTR

GtTR relies on use of a reference sample with accurate TR genotypes. This could be obtained from the long read capture sequencing of a reference sample, however, the drawback of this approach is that long-read sequencing would have to be obtained for each new TR target panel. An alternative to this approach is to use a sample which has been fully sequenced using long-read sequencing. Conveniently, Pendleton *et. al*. (2015) recently released a sample (NA12878) which has been sequenced to over 45X depth using PacBio [27].

VNTRtyper was applied to this sequencing data to calculate the number of repeats in our targeted TR regions. PacBio WGS data on NA12878 sample had sufficient coverage on 122 targeted TR regions to determine the genotype (Additional File 2 - Supplementary Table 4). The genotype estimates by VNTRtyper were compared to the genotypes determined by PCR sizing analysis for nine targets (Figure 5a and 5b). The correlation values between VNTRtyper and PCR sizing analysis indicate that the genotype predictions on PacBio WGS data by VNTRtyper were comparable to the accuracy of PCR results. On average, the VNTRTyper result was 13% different from the PCR sizing (Table 1) and this was mainly due to low coverage of PacBio WGS data.

**Figure 5:**
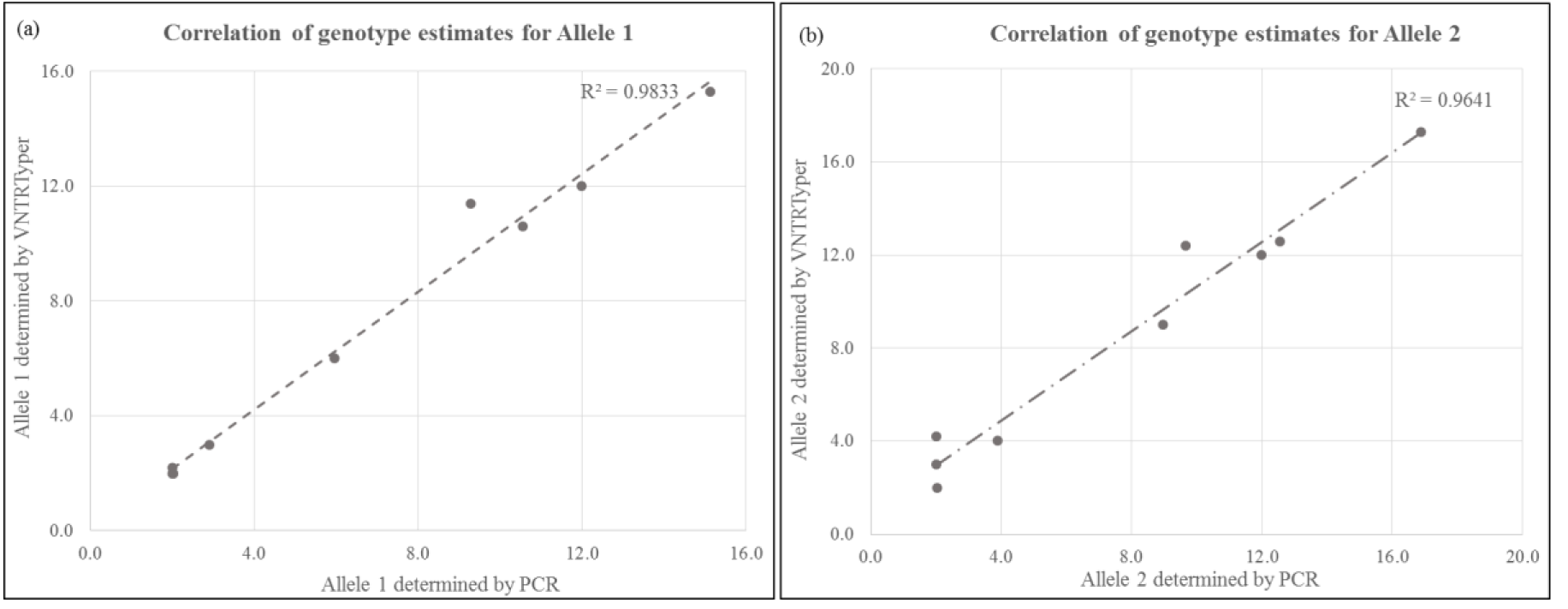
Comparison of allele calls between VNTRtyper on PacBio WGS and PCR sizing analysis on NA12878 sample for (a) allele 1 and (b) allele 2.

### Genotyping VNTRs using short-read Illumina targeted capture sequencing data

All of the targeted VNTRs were captured successfully by the targeted capture sequencing method and approximately 90% of the targets have greater than 100X coverage (Figure 6a). The sequence coverage was not affected by the GC content of the repeat sequences (Figure 6b) or the length of the repeat unit.

**Figure 6:**
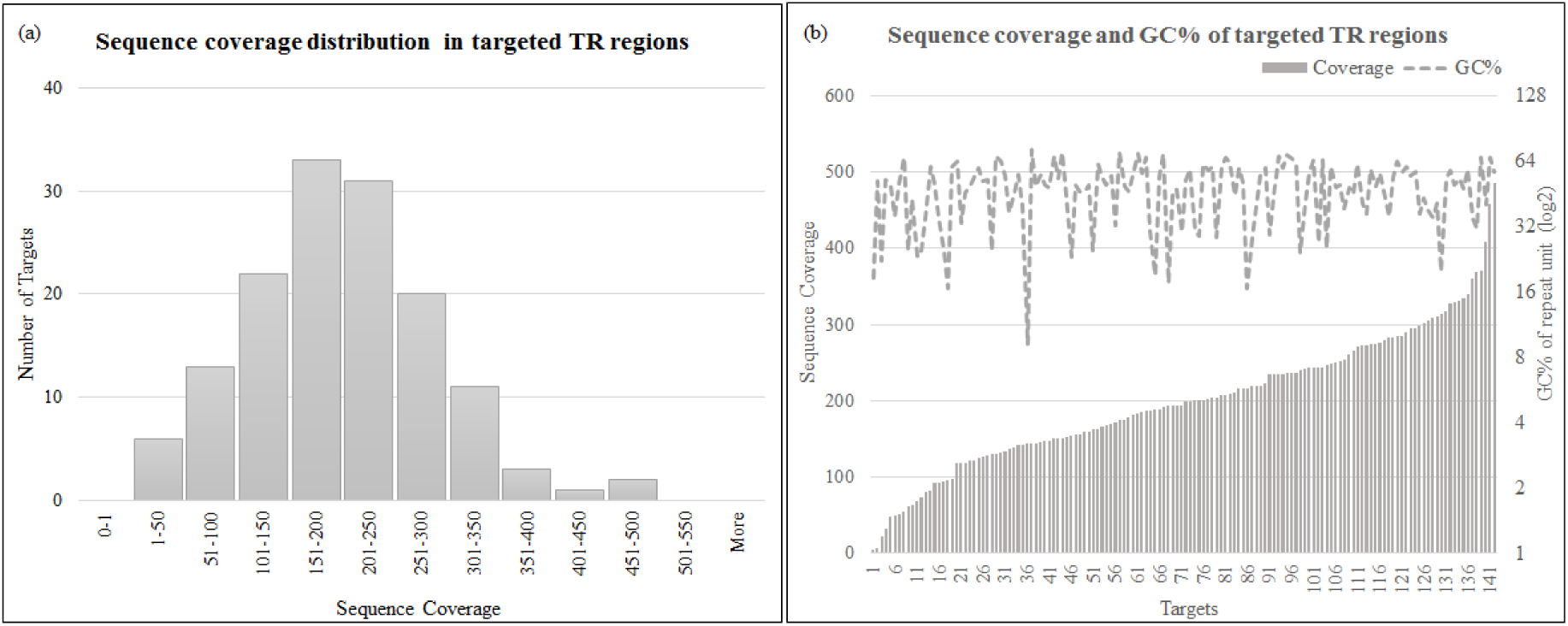
Coverage plots for Illumina targeted capture sequencing (a) average sequence coverage distribution in targeted TR regions (b) average sequence coverage distribution per target and GC% of the repeat unit.

The GtTR algorithm was applied to Illumina targeted sequencing data to determine the genotypes of the targeted VNTR regions. NA12878 PacBio WGS data (Additional File 2 - Supplementary Table 4) was used to calculate reference genotypes for 122 out of 142 targets; the remainder did not have sufficient depth to calculate accurate genotypes (see Methods). Two technical replicates were included for Illumina targeted sequencing of NA12878 sample. One of these replicates was used as reference for RCN calculations and the other was used as a test sample (see methods).

The genotype estimates by GtTR were compared to the average of two alleles determined by PCR on nine targets for all 7 samples (Table 1). Capillary electrophoresis plots for the PCR sizing analysis is provided in Additional file 1. The correlation values between genotype calls estimated from Illumina targeted sequencing and PCR sizing range from 0.9738 to 0.9930.

Genotypes estimates from GtTR using the Pacbio reference on the 9 loci with PCR sizing analysis were 68% accurate (CI: 55%-79%) using 95% HPD intervals (Figure 7a). However, if we restrict to the 6 loci for which the VNTRTyper estimate from the Pacbio reference was concordant with the PCR sizing result, the accuracy was 81% (CI: 65.9%-91.4%) (Figure 7b). If we had accurate reference genotypes for all 9 loci (i.e. PCR sizing estimate), then the accuracy would be 87.3% (CI 76.5% - 94.3%) (Figure 7c). This demonstrates the importance of obtaining highly accurate reference genotypes.

**Figure 7:**
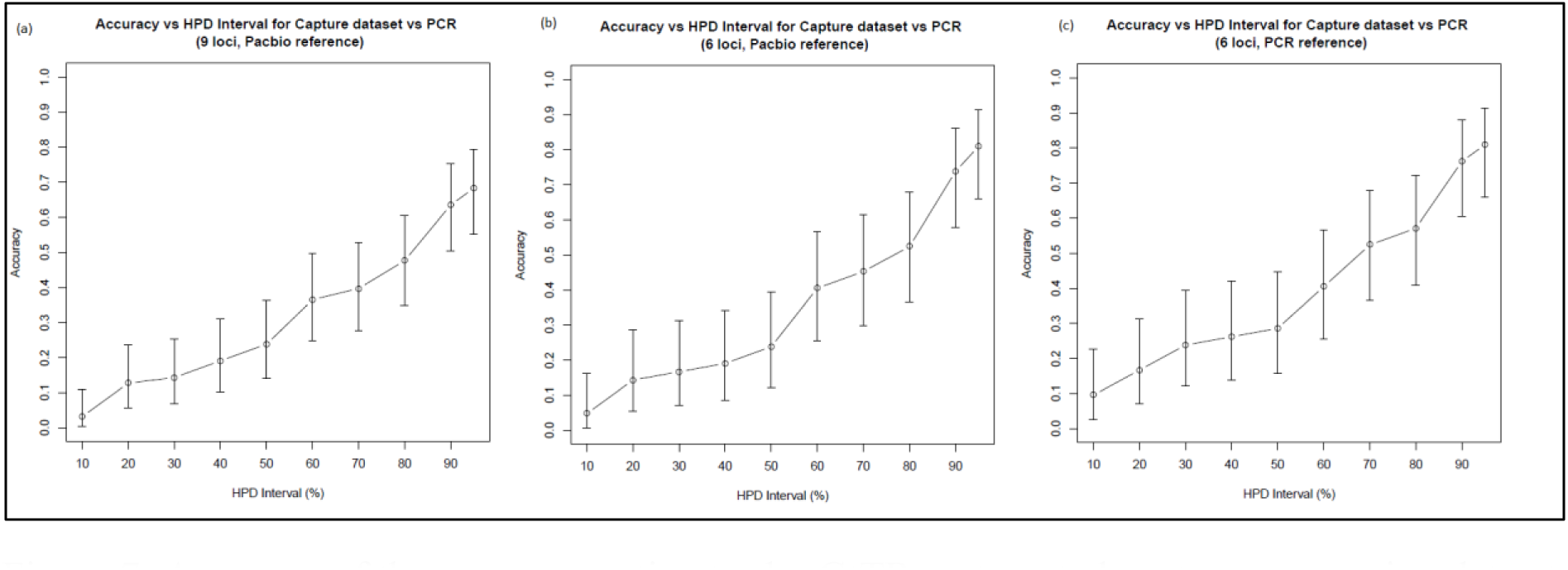
Accuracy of the genotype estimates by GtTR on targeted capture sequencing data at varying high posterior density intervals for (a) 9 targets validated by PCR sizing analysis using genotypes from PacBio sequence data as a reference genotype (b) 6 targets for which the Pacbio genotype estimates were concordant with the PCR sizing analysis using genotypes from PacBio sequence data as a reference genotype (c) 9 targets validated by PCR sizing analysis using genotypes from PCR analysis as a reference genotype. Error bars represent 95% binomial confidence intervals.

The median relative half-length of 95% HPD interval was 12.1% (across 122 loci and 7 samples with median depth of 395X). (Figure 8a). Amongst the 3 loci with depth greater than 800X the median relative half-length was 8.1% (Figure 8b), demonstrating the influence of read depth on the resolution of the estimates.

**Figure 8:**
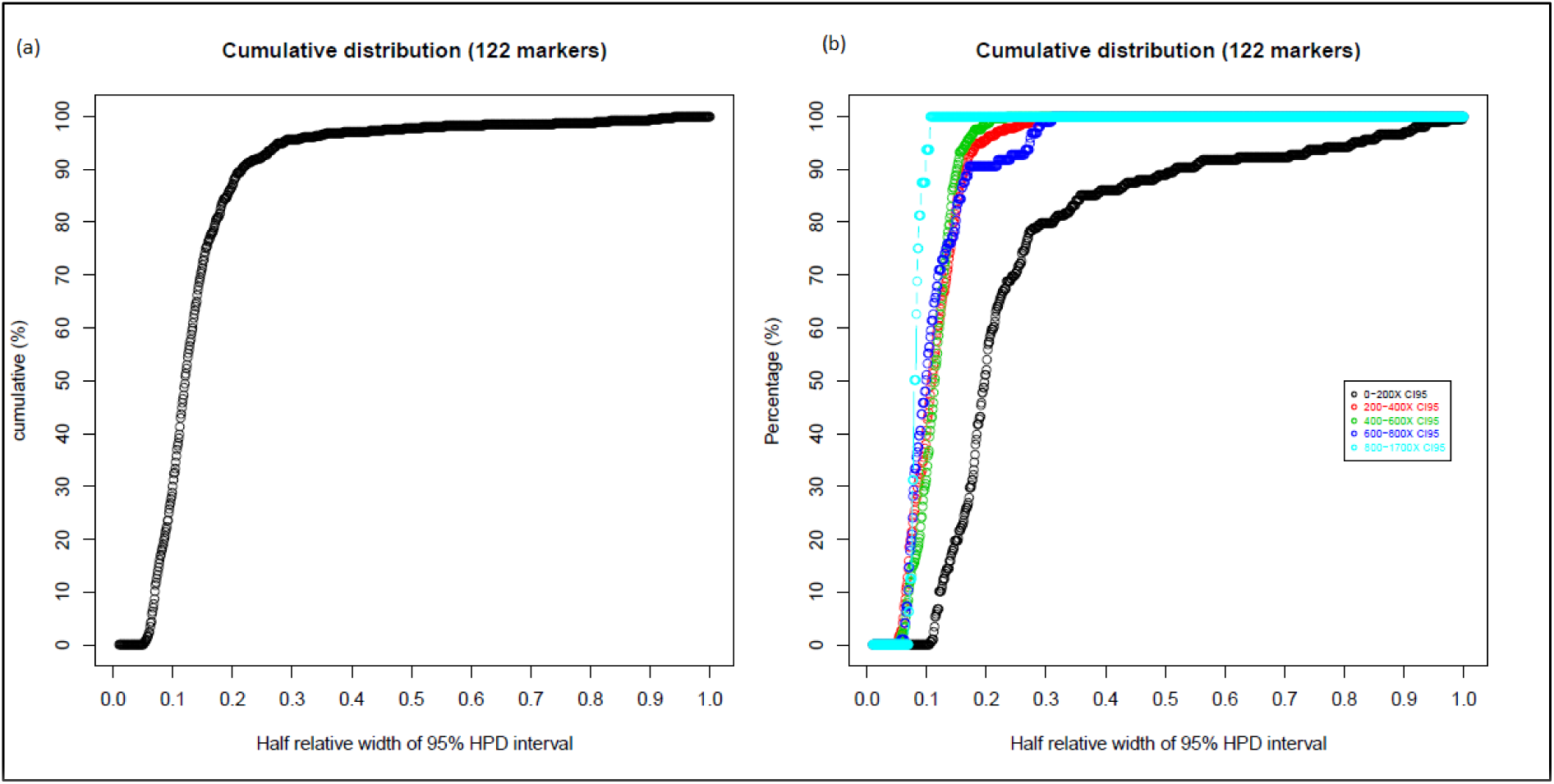
(a) Overall cumulative distribution of half relative width of 95% HPD interval in genotype estimates in targeted capture sequencing (b) Cumulative distribution of half relative width of 95% HPD interval in genotype estimates at varying sequence coverage.

### Genotyping VNTRs using short-read Illumina whole genome sequencing data

We downloaded 30X coverage Illumina WGS data on CEPH Pedigree 1463 for 17 samples including all 7 samples which were included in this study. Additionally we also downloaded 200X coverage sequencing data for 3 samples, including NA12877 and NA12878 samples included in this study. GtTR algorithm was applied to this WGS data set and NA12878 sample with 200X coverage was used as the reference sample for RCN analysis. Genotype estimates by GtTR on Illumina WGS data were compared with PCR sizing analysis for all 7 samples included in this study (Table 2). Comparison between genotype estimates by GtTR on Illumina WGS data and PCR sizing analysis for all Illumina WGS samples are provided in Supplementary Table 5 (Additional File 2).

Genotype estimates on Illumina WGS for the 9 targets validated by PCR were 89.6% accurate (CI:83.4% to 94.1%) and 83.3% (CI:58.5% to 96.4%) accurate using 95% HPD intervals at 30X coverage (using 17 samples) and 200X coverage (using 3 samples) respectively (Figure 9a). However, if we use the PCR genotypes as reference for the 9 targets, then these increase to 92.4% (CI: 86.7% - 96.1%) for 30X and 94.4% (CI: 72.7% - 99.9%) for 200X (Figure 9b). As before, this underscores the importance of obtaining accurate reference genotype calls. The median half relative width of 95% HPD interval (across 122 loci) was 25.0% and 14.8% for 30X and 200X coverage samples respectively (Figure 9c). This decreased to 11.7% amongst 7 loci in 200X coverage data which had greater than 400X coverage (Figure 9d).

**Figure 9:**
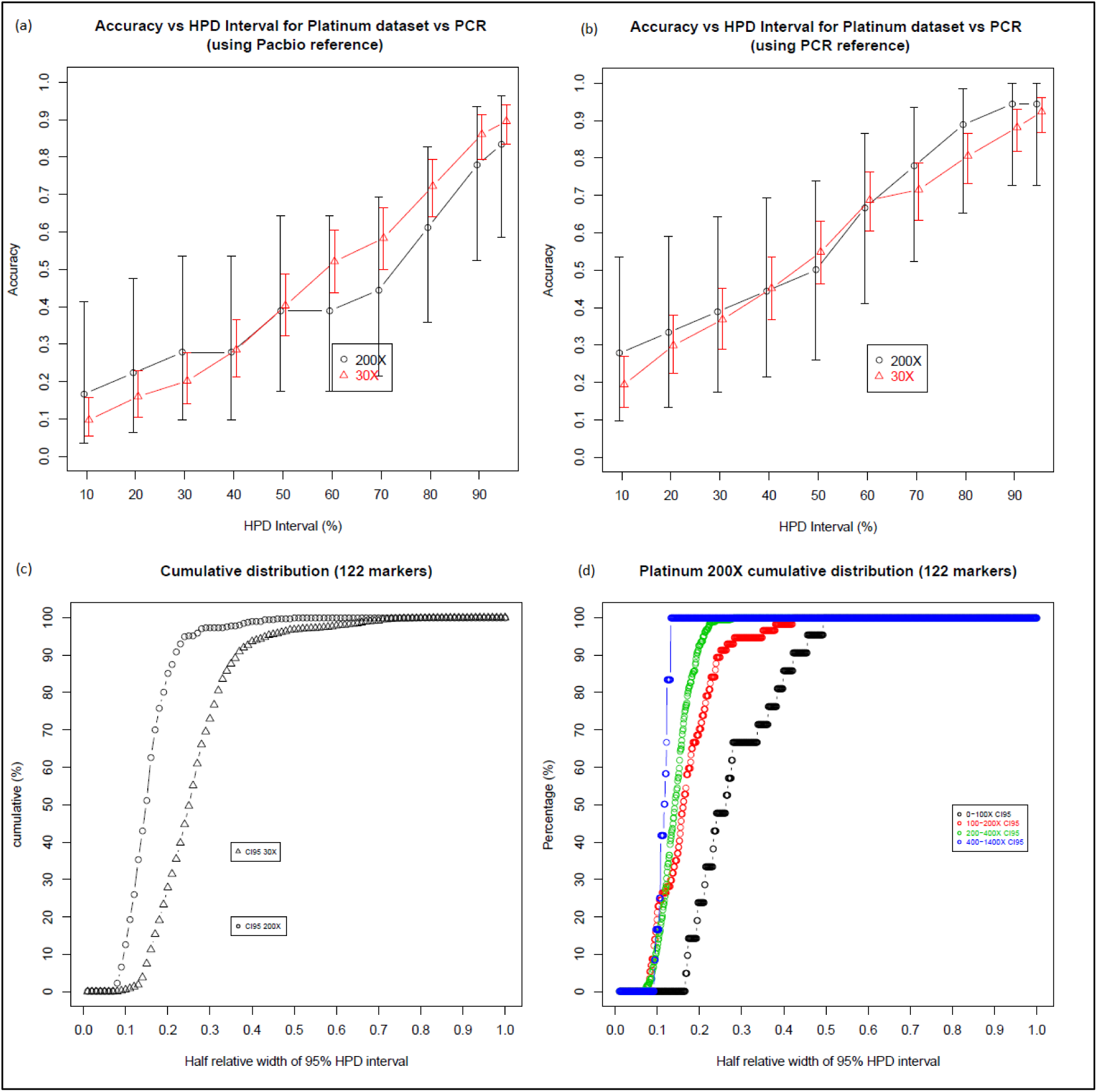
Accuracy of the genotype estimates by GtTR on Illumina WGS data at varying HPD intervals for 9 targets validated by PCR sizing analysis (a) using genotypes from PacBio sequence data as a reference genotype and (b) using genotypes from PCR sizing as a reference genotype. Error bars represent 95% binomial confidence intervals. (c) Cumulative distribution of half relative width of 95% HPD interval in genotype estimates in Illumina WGS 30X and 200X coverage (d) Cumulative distribution of half relative width of 95% HPD interval in genotype estimates at varying sequence coverage of Illumina 200X WGS.

Noticeably, the correlation values between Illumina targeted capture sequencing vs PCR sizing were higher than Illumina WGS vs PCR sizing (refer to Table 1 and Table 2), indicating that the high coverage targeted capture sequencing data improves the accuracy of genotype estimates. The correlation between genotype estimates on WGS and targeted capture sequencing data (Additional File 2 - Supplementary Table 6) range from 0.9340 to 0.9765. The low correlation values were likely due to the low accuracy in genotype estimates from low coverage (30X) WGS data. This is evident with NA12877 sample, where the correlation value improved from 0.9636 to 0.9800 (Additional File 2 - Supplementary Table 6), respectively for WGS 30X and 200X data.

## DISCUSSION

There are almost 1 million TRs in the human genome encompassing 4% of the entire genome including coding and regulatory regions [1]. Due to their unstable nature, TRs can lead to high rates of repeat variation and mutation in the genome [8]. Repeat variation in tandem repeats could exert functional consequences on adjacent genes [31]. Furthermore, variation in TRs are a major source of genomic variation between individuals and could possibly explain some of the phenotypic variation observed in complex diseases. However, the analysis of TRs are limited due to the lack of efficient high throughput analysis tools.

In this study, we present a novel high throughput targeted sequencing approach to genotype TRs. Our approach GtTR, uses short-read sequencing in combination with long-read characterized reference sample to genotype TRs in a cost-effective high throughput manner. Long reads, which span the entire repeat region and flanking region of the TRs enables accurate estimation of the number of repeats. Therefore the use of long-read sequencing data to determine the genotype of reference sample improves the accuracy of the genotype estimates by our ‘GtTR’ approach. Furthermore, the use of a global long-read WGS data as a reference sample data set (i.e. NA12878) eliminates the need to generate long read sequencing data on the reference sample for each new target panel. The reference sample only needs to be sequenced on a low cost targeted short-read sequencing method along with test samples for each new target panel.

The genotype estimates by GtTR on the targeted VNTR regions had comparable accuracy to PCR sizing analysis. Although we were only able to include nine regions for PCR validation due to the laborious nature of developing and validating PCR primers, the variation in repeat unit length, number of repeat units and sequence composition of the repeat in these nine regions provides a comprehensive representation of the entire targeted VNTR regions. We have also demonstrated our method GtTR on Illumina WGS data and the results reveal comparable accuracy to PCR sizing analysis. However, it was evident that the high sequencing coverage achieved from targeted sequencing provided an advantage for accurate genotype estimation in targeted regions.

One of the main drawbacks of our GtTR approach is that due to the use of read depth based analysis, the genotype estimates are an average of the 2 alleles instead of the exact estimates of 2 alleles as with long-read sequencing data. Although this might prevent the estimation of exact genotype, we believe this approach might still be applicable in GWAS. Difference in genotype estimates between test and control samples might be sufficient to identify TRs which might be associated with a complex disease. It is also worth noting that the genotype estimates from GtTR are dependent on a reference, therefore the genotypes are relative to reference sample and not the exact genotype. Thus, any errors in the reference sample will affect the estimates in the test samples.

There have been several studies on the use of targeted sequencing of STRs using short-read sequencing [20, 32-35]. However, the shorter read length of these technologies presents a challenge for genotyping longer TR regions. To our knowledge, our study is the first to successfully demonstrate targeted capture sequencing and genotyping of VNTRs up to 5Kb in length using short read sequencing. Our combination of long-read reference genotyping and short-read sequencing has enabled us to genotype difficult repetitive sequences and our approach has provided a cost-effective solution to genotype hundreds of TRs simultaneously in multiple samples.

Recently, a similar hybrid approach, MixTaR was published, which combines the high-quality of short-reads and the longer length of long-reads for tandem repeat detection [36]. However, this method requires the sample to be sequenced using both long-read and short-read sequencing methods to genotype TRs. Although, MixTaR provides an accurate genotype of 2 alleles, it is not feasible to apply this approach in population based genotyping studies. In contrast our approach uses a global reference sample, which eliminates the need to generate long read sequencing data for each sample, providing a cost effective option for population scale studies.

## CONCLUSIONS

In summary, we present a novel approach to enable genotyping longer TR regions using short-read sequencing. Using this method, we have successfully demonstrated the feasibility of targeted capture sequencing of repetitive sequences and genotyping VNTRs longer than the short-read sequence length. We believe our approach would provide a tool for large scale genome-wide population analysis to assess the impact of tandem repeat variability in complex traits.

## LIST OF ABBREVIATIONS

TR: Tandem repeat
STR: Short tandem repeat
VNTR: Variable number of tandem repeat
SNP: Single nucleotide polymorphism
CNV: Copy number variation
GWAS: Genome wide association studies
RFLP: Restriction fragment length polymorphism
PCR: Polymerase chain reaction
TRDB: Tandem Repeats Database
PCA: Principal component analysis
HMM: Hidden markov model
WGS: Whole genome sequencing
HPD: High Posterior Density
CI: Confidence Interval
HRW: Half Relative Width

## DECLARATIONS

Availability of data and materials: The sequencing datasets generated during the study are available in NCBI SRA repository, accession number SRP126797 https://www.ncbi.nlm.nih.gov/sra/SRP126797.

## Authors’ contributions

DG, MDC and LC conceived the study. DG and TD performed the experiments. WC performed the analysis for target selection. DG, MDC and LC performed the sequencing and statistical analysis. DG wrote the paper with input from the other authors.

## Acknowledgements

We would like to acknowledge Dr. Daniele Belluoccio for his assistance with target capture design process, Dr. Tim Bruxner and Angelika Christ for their support with sequencing.

## Competing interests

The authors declare that they have no competing interests.

## Funding

This study is supported by the funding from Australian National Health and Medical Research Council (APP1052303).

## Ethics approval and consent to participate

Not applicable

## Consent for publication

Not applicable

## ADDITIONAL FILES

Additional file 1: Supplementary information on analysis and capillary electrophoresis results of PCR sizing (PDF 1264 kb)

Additional file 2: Supplementary Tables S1-S5 (XLSX 295 kb)

